# Temperature regulation as a tool to program synthetic microbial community composition

**DOI:** 10.1101/2020.02.14.944090

**Authors:** Adam Krieger, Jiahao Zhang, Xiaoxia Nina Lin

## Abstract

Engineering of synthetic microbial communities is emerging as a powerful new paradigm for performing various industrially, medically, and environmentally important processes. To reach the fullest potential, however, this approach requires further development in many aspects, a key one being regulating the community composition. Here we leverage well established mechanisms in ecology which govern the relative abundance of multi-species ecosystems and develop a new tool for programming the composition of synthetic microbial communities. Using a simple model system consisting of two microorganisms *Escherichia coli* and *Pseudomonas putida*, which occupy different but partially overlapping thermal niches, we demonstrate that temperature regulation can be used to enable coexistence and program the community composition. We first investigate a constant temperature regime and show that different temperatures lead to different community compositions. Next, we invent a new cycling temperature regime and show that it can dynamically tune the microbial community, achieving a wide range of compositions depending on parameters that are readily manipulatable. Our work provides conclusive proof of concept that temperature regulation is a versatile and powerful tool capable of programming compositions of synthetic microbial communities.

## 1. Introduction

Microbial systems have been used for decades to perform functions with industrial, environmental, and medical relevance (Bouwer & Zehnder, 1993; Göddel et al., 1979; Houde, Kademi, & Leblanc, 2004). Traditionally, a single microbial species is cultured for a specific functionality. If necessary, genetic modifications are made to this organism to enable novel capabilities. As the field surrounding this paradigm has progressed, however, challenges faced by this monoculture approach have become increasingly apparent. Metabolic versatility and complexity in a single species is limited by factors such as metabolic burden, a lack of or limited intracellular compartmentalization, and energetic tradeoffs (Litchman, Edwards, & Klausmeier, 2015; Zhou, Qiao, Edgar, & Stephanopoulos, 2015). At the population level, monocultures are susceptible to invasion and subsequent population collapse by agents such as foreign bacteria, fungi, and bacteriophages. One approach to address these challenges is utilization of synthetic microbial communities as opposed to monocultures for biotechnology applications. Microbial communities enable division of labor and specialization within a population, decreasing metabolic burden and enabling reaction compartmentalization. More diverse microbial communities can also exhibit increased stability (Yurtsev, Conwill, & Gore, 2016) and resistance to invasion by foreign agents (Dirk et al., 2012), as well as increased biomass production (McGrady-Steed, Harris, & Morin, 1997) and nutrient utilization (Ptacnik et al., 2008) in certain contexts.

While the potential advantages offered by a microbial community over a monoculture are clear, community based approaches face challenges of their own. One of the most basic challenges is control of community composition. Community composition can have different meanings in different ecological contexts, but here is used to refer specifically to the relative abundance of individual species within the community. In a microbial community in which each species exhibits a specific and unique function within the community, the metabolic functionality of the community as a whole can depend greatly on the relative abundance of each species within the community (Hsu et al., 2019; Scholz, Graves, Minty, & Lin, 2017; H. Zhang, Pereira, Li, & Stephanopoulos, 2015). Despite significant advances in many related research topics, progress in control of synthetic microbial community compositions has been slow, likely due at least in part to a dearth of tools capable of dynamically modulating relative species abundance. The most commonly used technique is manipulation of inoculation ratio (Minty et al., 2013; Shong, Rafael, Diaz, & Collins, n.d.; Strickland, Lauber, Fierer, & Bradford, 2009), but this approach is not capable of dynamic control and due to differences between intrinsic growth rates and interspecific interactions between community residents the community composition often quickly shifts away from this initial condition.

While synthetic biology researchers are starting to develop new approaches to address the aforementioned challenges, mechanisms that enable species coexistence and modulate relative abundance over time in complex naturally occurring communities have been a major theme in ecological study for almost 100 years. The current state of the field proposes that all mechanisms that maintain and modulate species diversity operate through one of two broad classes, equalizing mechanisms and stabilizing mechanisms (Chesson, 2000). Equalizing mechanisms lessen the relative fitness differences between species (which can be more simply conceptualized as growth rates, although in truth this is a slight oversimplification), and stabilizing mechanisms increase the niche differences between species, which is to say they decrease the extent to which species compete directly with each other (Chesson, 2000). According to Chesson’s theory, equalizing mechanisms can support coexistence by decreasing the difference between growth rates, but stabilizing mechanisms are required for coexistence. This makes intuitive sense in that, if there is a difference between the growth rates of two species, no matter how small, eventually that difference will lead to competitive exclusion of the slower growing species in the absence of additional compensatory mechanisms. Therefore, any environmental variables which differentially affect growth rates or niche partitioning would be expected to influence the ability of species to coexistence and/or community composition.

In this work we have applied concepts from the ecological study of the mechanisms enabling maintenance of species coexistence to a synthetic microbial community in order to develop tools that are capable of dynamically modulating relative species abundance. The modality we have implemented is temperature regulation. Using a synthetic community consisting of two model microorganisms *Escherichia coli* and *Pseudomonas putida*, which occupy different but partially overlapping thermal niches, we demonstrate that temperature regulation is a tool that can be used to enable coexistence and program the community composition. Specifically, we develop two temperature mediated regimes. First, using a constant temperature for the co-culture, we are able to identify a temperature window that allows co-existence of the two microbes, albeit exhibiting only a small range of achievable compositions. Next, we develop a novel regime of cycling between a high temperature favoring *E. coli* and a low one favoring *P. putida*, which enables co-existence of the two microorganisms in a highly tunable manner.

## 2. Materials and Methods

### 2.1 Microbial strains and cultivation

*E. coli K12 substr. MG1655* constitutively expressing a chromosomally integrated YFP construct was generated by P1 transduction as described by Thomason et al.(Thomason, Costantino, & Court, 2007) Strain Y2 from Kerner et al.(Kerner, Park, Williams, & Lin, 2012) was used as the source of the YFP cassette. The *P. putida KT2440* strain constitutively expressing mCherry was a generous gift from Dr. Esteban Martinez-Garcia of the Victor de Lorenzo lab. Addition of an mCherry cassette to *P. putida KT2440* was performed via Tn7 transposon assisted cloning (Koch, Jensen, & Nybroe, 2001; Martínez-García, Calles, Arévalo-Rodríguez, & De Lorenzo, 2011).

Unless otherwise noted, all cultures were grown in minimal M9 media (200 mL 5x M9 salts [34 g/L Na_2_HPO_4_ anhydrous, 15 g/L KH_2_PO_4_, 2.5 g/L NaCl, 5 g/L NH_4_Cl], 20 mL 20% glucose, 2 mL 1M MgSO_4_, 100 uL 1M CaCl_2_, 780 mL sterile deionized H_2_O; autoclave all components separately then mix under sterile conditions).

### 2.2 Visualization of strains via confocal microscopy

50 uL of log phase bi-culture was pipetted onto a slide and a coverslip was placed on top. Confocal fluorescence microscopy was performed using an upright Olympus FV1200 confocal microscope equipped with a 60x objective and 405, 488, 515, 543, and 635 lasers. Images were exported and analyzed with ImageJ.

### 2.3 Quantification of maximum specific growth rate in co-cultures

*E. coli K12 substr. MG1655-YFP* and *P. putida KT2440-mCh* were seeded from -80 °C cryostocks and grown in monoculture overnight to stationary phase in 14 mL Corning Falcon^®^ test tubes (polypropylene test tube, round bottom, 17×100mm, 14mL, graduated, with clear snap cap, Sterile, 25 per Pack) in a 2 mL volume of M9 minimal media. Overnight cultures were diluted 1:100 into fresh M9 media and grown into exponential growth phase (OD_600_ ∼0.4-0.6). The cell density of the two cultures of exponential phase cells were normalized to each other using OD_600_ and then diluted 1:100 into fresh M9 media in a Greiner Bio-one CELLSTAR™ 96 well μClear flat bottomed microplate with a 200 uL final volume. Strains were grown in monoculture and co-culture in triplicate (2 uL of the desired strain was added for monocultures, and 1 uL each of both strains was added for co-cultures to 198 uL M9 media). These nine wells were surrounded by wells filled with 200 uL sterile deionized H_2_O to inhibit evaporation from experimental wells. The lid was treated with a mixture of 20% ethanol + 0.5% Triton X-100 to avoid condensation formation on the lid (pour enough mixture to completely cover the bottom of the lid, let sit 5 minutes, pour off and let air dry). The lid was taped on using Fisherbrand™ labeling tape. The plate was incubated in a Biotek Synergy H1 platereader for 24 hours with plate reads every 10 minutes and continuous orbital shaking at 282 cpm. At each timepoint, reads at 600 nm wavelength, Excitation: 510 nm Emission: 540 nm, and Excitation: 585 nm Emission: 620 nm were taken of each well (these wavelengths were empirically determined to maximize the specific signal and minimize crosstalk between the two channels for our constructs, media, and platereader; data not shown). The maximum specific growth rates from the co-culture growth curves were calculated by fitting the early exponential growth phase to an exponential function using Microsoft Excel.

### 2.4 Constant temperature regime: culture growth and community composition quantification

*E. coli K12 substr. MG1655-YFP* and *P. putida KT2440-mCh* were seeded from -80 °C cryostocks and grown in monoculture overnight to stationary phase in 14 mL Corning Falcon^®^ test tubes (polypropylene test tube, round bottom, 17×100mm, 14mL, graduated, with clear snap cap, Sterile, 25 per Pack) in a 2 mL volume of M9 minimal media. Overnight cultures were diluted 1:100 into fresh M9 media and grown into exponential growth phase (OD600 ∼0.4-0.6). The cell density of the two cultures of exponential phase cells were normalized to each other using OD_600_ and then diluted into 14 mL Corning Falcon^®^ test tubes with 2 mL fresh M9 at three different inoculation ratios; 1:10, 1:1, and 10:1 *E. coli*: *P. putida* (2 uL *E. coli* + 18 uL *P. putida*, 10 uL *E. coli* + 10 uL *P. putida*, and 18 uL *E. coli* + 2 uL *P. putida*). Each inoculation ratio was performed in triplicate producing a total of nine cultures. Cultures were incubated in a Lab-Line Instruments Model No. 3528 incubator with a New Brunswick Scientific C1 platform shaker placed inside for shaking set at a speed of 50 (no units are indicated on the shaker). The temperature was monitored throughout the lifetime of each experiment using an Omega OM-91 portable temperature data logger (see Supplemental Information). Each culture was passaged 1:100 into 2 mL fresh M9 media in 14 mL Corning Falcon^®^ test tubes twice daily (8 hour and 16 hour growth periods). At each passaging time point, each culture was vortexed for 5 seconds and a 1 uL sample was taken for quantification of community composition via flow cytometry. Passaging was performed after vortexing.

### 2.5 Cycling temperature regime: culture growth and community composition quantification

*E. coli K12 substr. MG1655-YFP* and *P. putida KT2440-mCh* were seeded from -80 °C cryostocks and grown in monoculture overnight to stationary phase in 14 mL Corning Falcon^®^ test tubes (polypropylene test tube, round bottom, 17×100mm, 14mL, graduated, with clear snap cap, Sterile, 25 per Pack) in a 2 mL volume of M9 minimal media. Overnight cultures were diluted 1:100 into fresh M9 media and grown into exponential growth phase (OD_600_ ∼0.4-0.6). The cell density of the two cultures of exponential phase cells were normalized to each other using OD_600_ and then diluted into a 14 mL Corning Falcon^®^ test tube with 2 mL fresh M9 at a 1:1 inoculation ratio (10 uL *E. coli* + 10 uL *P. putida*). For the long-time interval cycling temperature program experiments, the culture was incubated at 37 °C for 16 hours in a New Brunswick Scientific Excella E24 Incubator Shaker with 225 rpm shaking, and at 31 °C for 8 hours in a Lab-Line Instruments Model No. 3528 with 225 rpm shaking. For the short-time interval cycling temperature experiments, a culture was prepared as above and incubated for approximately 72 hours to allow adaptation to culture conditions. The culture was then incubated at 27 °C until the community composition reached a 1:1 ratio, and then was incubated at the indicated temperatures for the indicated times. After each growth period, cultures were vortexed for 5 seconds and a 1 uL sample was taken for quantification of community composition by flow cytometry, and then the culture was passaged 1:100 into 2 mL of fresh M9 media in a new 14 mL Corning Falcon^®^ test tube.

### 2.6 Quantification of community composition via flow cytometry

Samples taken from co-cultures were diluted to ∼10^6^ cell/mL in 1x PBS (137 mM NaCl, 2.7 mM KCl, 10 mM Na_2_HPO_4_, 1.8 mM KH_2_PO_4_; pH 7.4) and run on an Applied Biosystems Attune acoustic focusing cytometer. 300 uL of sample was acquired by the device for each run and 10,000 events were recorded. The instrument configuration was as follows: (Threshold (x1000): FSC: 10; SSC: 10; BL1: 10; BL2: 10; BL3: 10; BL4: 10; RL1: 10; RL2:10, Voltage (mV): FSC: 3,000; SSC: 3,500; BL1: 2,400; BL2: 1,800; BL3: 2,550; BL4: 2,550; RL1: 2,900; RL2: 1,950). Because this machine lacks a laser with proper wavelengths for detection of our mCherry construct, yellow fluorescence positive cells were classified as *E. coli K12 substr. MG1655-YFP* and yellow fluorescence negative cells were classified as *P. putida KT2440-mCh* (Fig. 3 A).

### 2.7 Modeling of population dynamics under the cycling temperature regime

For the cycling temperature regime, consider each cycle consists of a time interval *t*^*H*^ at a high temperature favoring *E. coli* and then another time interval *t*^*L*^ at a low temperature favoring *P. putida*. Two types of models were developed. In the first one, simple exponential growth is assumed.

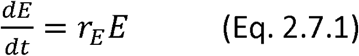

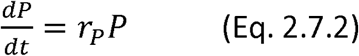

where variables *E* and *P* are *E. coli* and *P. putida* cell densities respectively; *r*_*E*_ and *r*_*P*_, their specific growth rates respectively, are model parameters. We further denote 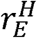 and 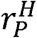 as the specific grow rates of *E. coli* and *P. putida* respectively at the high temperature; 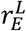 and 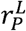 the specific grow rates of *E. coli* and *P. putida* respectively at the low temperature. Note that 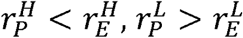, and we define 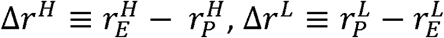.

The above ODE’s have simple analytical solutions and the two cell densities can be expressed as explicit functions of time:

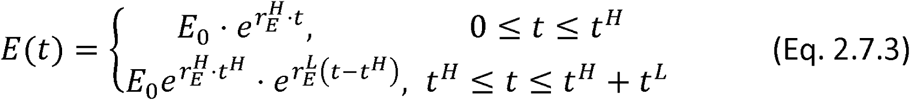

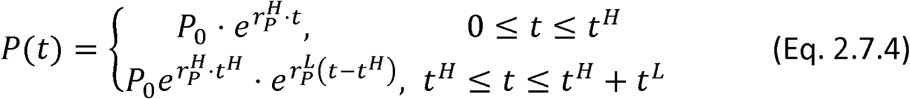

where *E*_0_ and *P*_0_ are the *E. coli* and *P. putida* cell densities respectively at the beginning of the cycle.

To maintain the bi-culture stably, the composition at the end of each cycle needs to return to its value at the beginning of the cycle. Therefore, we require the following:

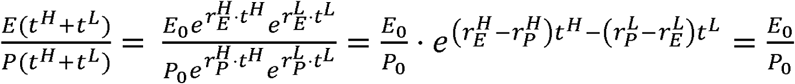

It follows that:

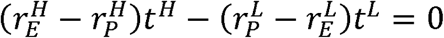

Therefore, we have the relationship (Eq. 3.4.1):

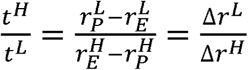

The ratio of the two species averaged over the whole cycle can be calculated as follows:

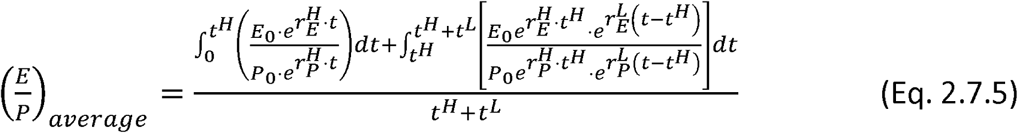

Simplifications of this expression, including utilization of the relationship above between time intervals and growth rate differences required for maintaining the bi-culture, ultimately lead to the following (Eq. 3.4.2):

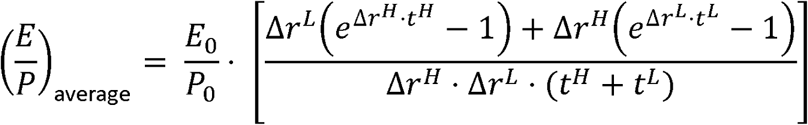

We also developed a second type of mathematical model where logistic growth is assumed.

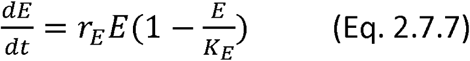

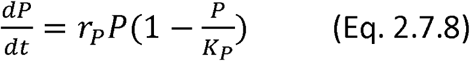

where *r*_*E*_ and *r*_*P*_ are the maximum specific growth rates of *E. coli* and *P. putida* respectively; *K*_*E*_ and *K*_*P*_ are the corresponding carrying capacities. A Mathlab script (see Supplemental information) was developed to simulate repeated cycles of cell growth at a high temperature and then low temperature using these two un-coupled ordinary differential equations.

## 3. Results

### 3.1 Overall approach

Our objective is to develop new tools for regulating synthetic microbial community compositions, by leveraging temperature-dependent growths of distinct desired community members. As illustrated in Fig. 1 for a simple community consisting of two members exhibiting different growth phenotypes at various temperatures, we propose two temperature mediated regimes for achieving co-existence and regulating community compositions. In the first regime (Fig. 1A), a constant temperature is used for growing the bi-culture. It is expected that if a low temperature is selected, the microbe favored at the temperature (i.e. exhibiting a higher growth rate) would dominate; and vice versa at a high temperature. We hypothesize, however, for a small range of intermediate temperatures, the two microbes could co-exist. In the second regime (Fig. 1B), the bi-culture is cycled between a low temperature and a high one, which favor each of the two microbes respectively, and co-existence of the two microbes are maintained over an extended period of time. Operating parameters include the high and low temperatures, and the time durations the bi-culture spends at each temperature. By manipulating these parameters, one would be able to regulate the average community composition and its variation.

**Figure 1.**
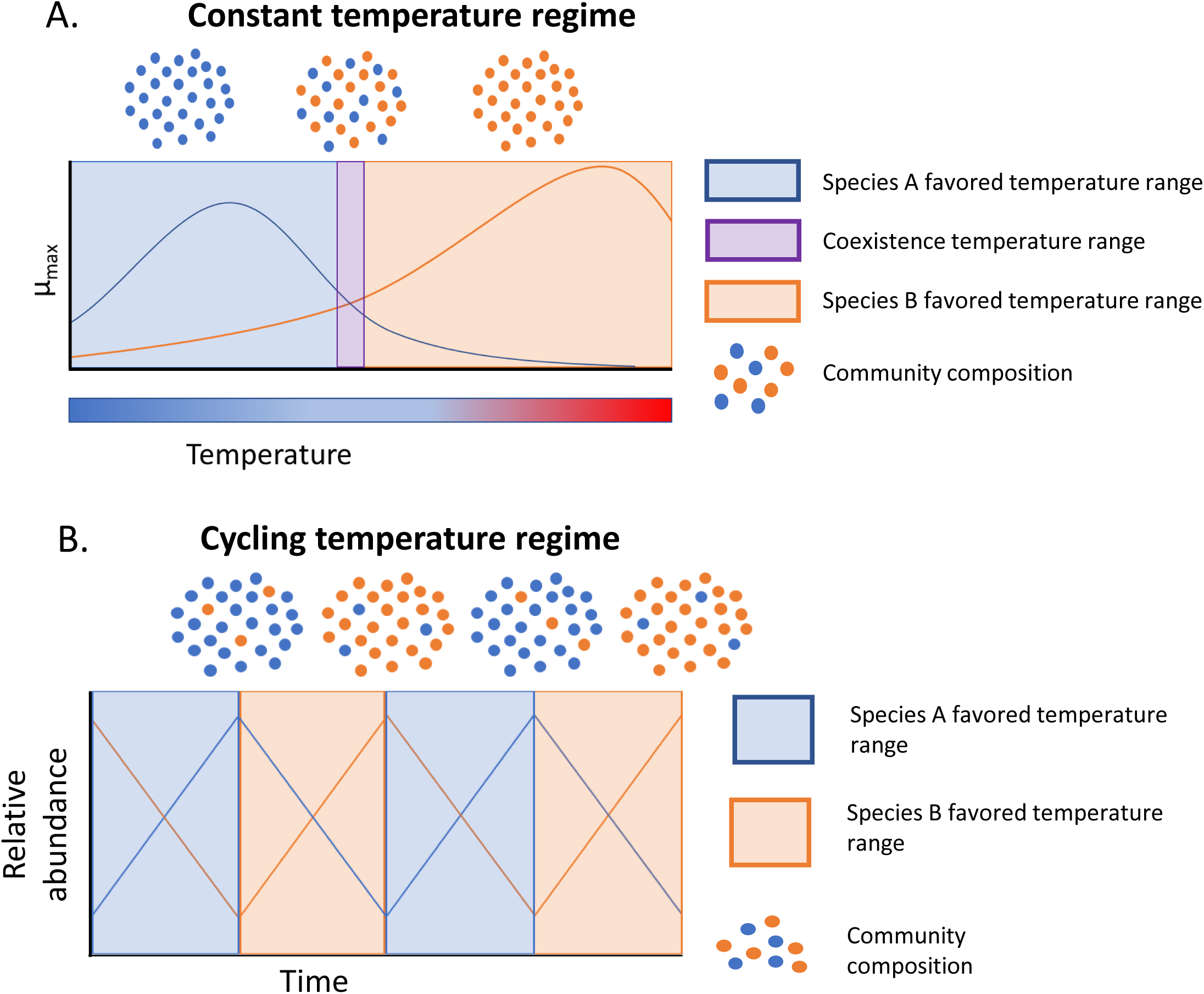
Two approaches for rationally regulating synthetic microbial community compositions. **A)** Under what we refer to as a constant temperature regime, a synthetic bi-culture is kept at a constant temperature throughout the duration of the culture lifetime. The selected temperature determines whether a monoculture persists or co-existence of the two microbes can occur. **B**) Under a cycling temperature regime a bi-culture is grown in repeated cycles of high and low temperatures, enabling co-existence of the two microbes over an extended period of time. Both the temperatures and the time intervals spent at each temperature can be manipulated to achieve desired outcomes.

### 3.2 Design and implementation of a model microbial bi-culture

To explore the temperature regulation regimes proposed above, we first established a model synthetic microbial system consisting of two well-characterized bacterial species. One is the soil bacterium *Pseudomonas putida KT2440* (Belda et al., 2016). *P. putida* is found throughout the natural environment, predominantly in soil and plant rhizospheres (Belda et al., 2016; Thomas, Koh, Bankowski, & Seto, 2013). This bacterium is a mesophilic organism, growing optimally at relatively moderate temperatures (Munna, M. S., Zeba, Z., and Noor, 2015), and has an extremely versatile metabolism which has earned it a reputation as an excellent bioremediation agent (Belda et al., 2016). In this study we use a strain of *P. putida* which has been tagged with a constitutively expressed chromosomal copy of mCherry fluorescent protein (mCh) (Rochat, Péchy-tarr, Baehler, Maurhofer, & Keel, 2010). The second bacterial species in our system is *E. coli K12 MG1655* (Blattner et al., 1997). One of the most commonly used model organisms in microbiology, *E. coli* can be either commensal or pathogenic gut colonizers of humans and other mammals (depending on the strain), growing optimally at relatively higher temperatures compared to *P. putida*. In this study we used a strain of *E. coli K12 MG1655* which has been tagged with a constitutively expressed chromosomal copy of yellow fluorescent protein (YFP). Microbial species have naturally evolved to modulate their growth rate in response to temperature, often with optimal growth occurring within a small range of temperature. We consider this pair of bacteria as a model system of microbial communities composed of members occupying different but partially overlapping thermal niches. They can be grown together in a bi-culture in minimal M9 medium.

We verified expression of the fluorescent proteins in each strain with confocal microscopy (Supplemental Fig. 2 A) and flow cytometry (Supplemental Fig. 2 B). We then quantified the growth phenotype of each species in co-culture at various temperatures. Particularly, we determined the maximum specific growth rates at different temperatures (Fig. 2, referred to as temperature profiles hereafter), using growth curves derived from population level fluorescence from each species in bi-cultures. The temperature profiles indicate that for each species there exists a temperature range in which the species would be expected to have a growth advantage over the other (when the temperature is less than ∼35.5 °C, *P. putida* μ_max_ > *E. coli* μ_max_ and when the temperature is greater than ∼35.5 °C then *P. putida* μ_max_ < *E. coli* μ_max_).

**Figure 2.**
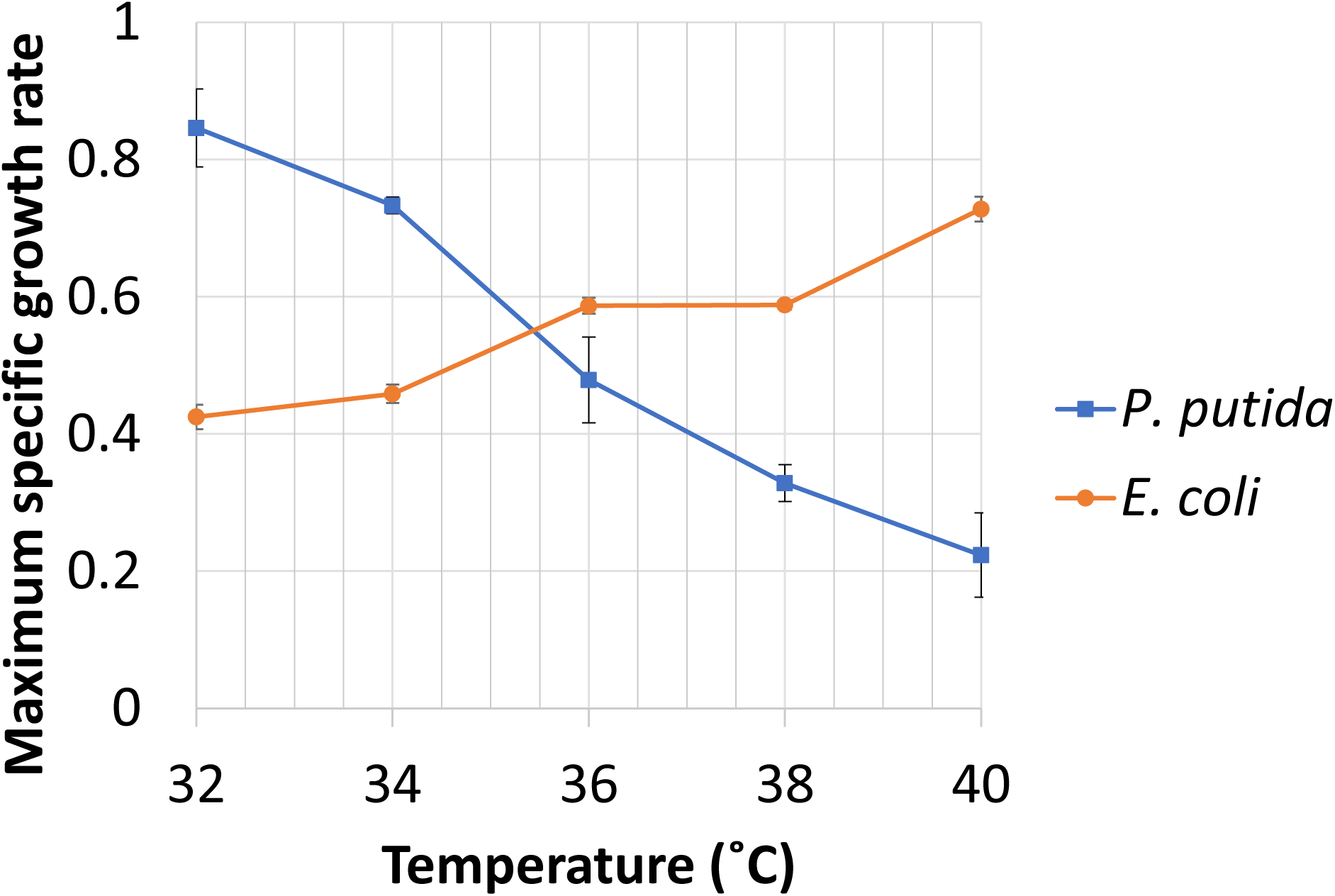
Maximum specific growth rates of each bacterium in co-cultures with different temperature profiles. *E. coli K12 substr. MG1655* grows optimally at relatively higher temperature and *Pseudomonas putida KT2440* grows optimally at relatively lower temperatures.

### 3.3 Effects on community composition of a constant temperature program

We first explore the constant temperature regime by investigating the community population dynamics over time at various selected temperatures. In this series of experiments, the temperature was maintained at a constant value (±0.2 °C, Supplemental Fig. 3) throughout the entire course of an experiment, and co-cultures were diluted 1:100 into fresh media twice daily (9 h and 15 h growth periods). Each experiment was conducted for times ranging from 60 to 400 hours depending upon the observed dynamics. Community composition was quantified via flow cytometry with a measurement taken at each dilution time point. The dynamics of the bi-culture is shown in Fig. 3A for three representative temperatures, 35.5 °C, 36.5 °C, and 39.5 °C; the full set of data for 10 temperatures in the range of 32-39.5 °C are provided in detail in Supplemental Fig. 1 and summarized in Fig. 3B. It was found that at 35.5 °C and lower, the community was dominated by *P. putida* when the culture composition reached equilibrium (which is defined herein as changing no more than ±15% over at least 3 consecutive time points), regardless of imposed initial ratios of the two bacteria (for example, see Fig. 3A top row). On the other hand, at temperatures above ∼39 °C the community was dominated by *E. coli* (for example, see Fig. 3A bottom row). Interestingly, at temperatures between 35.5 °C and ∼39 °C, coexistence of the two bacteria was observed (for example, see Fig. 3A middle row). It is worth noting that the community composition at equilibrium of this bi-culture depends on the temperature in a highly nonlinear manner. Specifically, the equilibrium composition changed very gradually in the relatively wide temperature window of 35.7-39.1 °C; whereas this property changed drastically over a much narrower temperature window of 35.5-35.7 °C. These differences have direct implications for practical applications where a constant temperature is employed. Given the resolution of temperature control (e.g. ±0.2 °C in our laboratory set-up), it is practically impossible to achieve a specific desired community composition in the small temperature window where the composition is highly sensitive to fluctuations in temperature. In contrast, in the temperature window where composition changes are milder, it is indeed possible to select a temperature to maintain the desired community composition. However, it is important to note that the range of community compositions achievable in this temperature window (∼70-95% *E. coli* at 36.1-39.1 °C) is rather limited.

**Figure 3.**
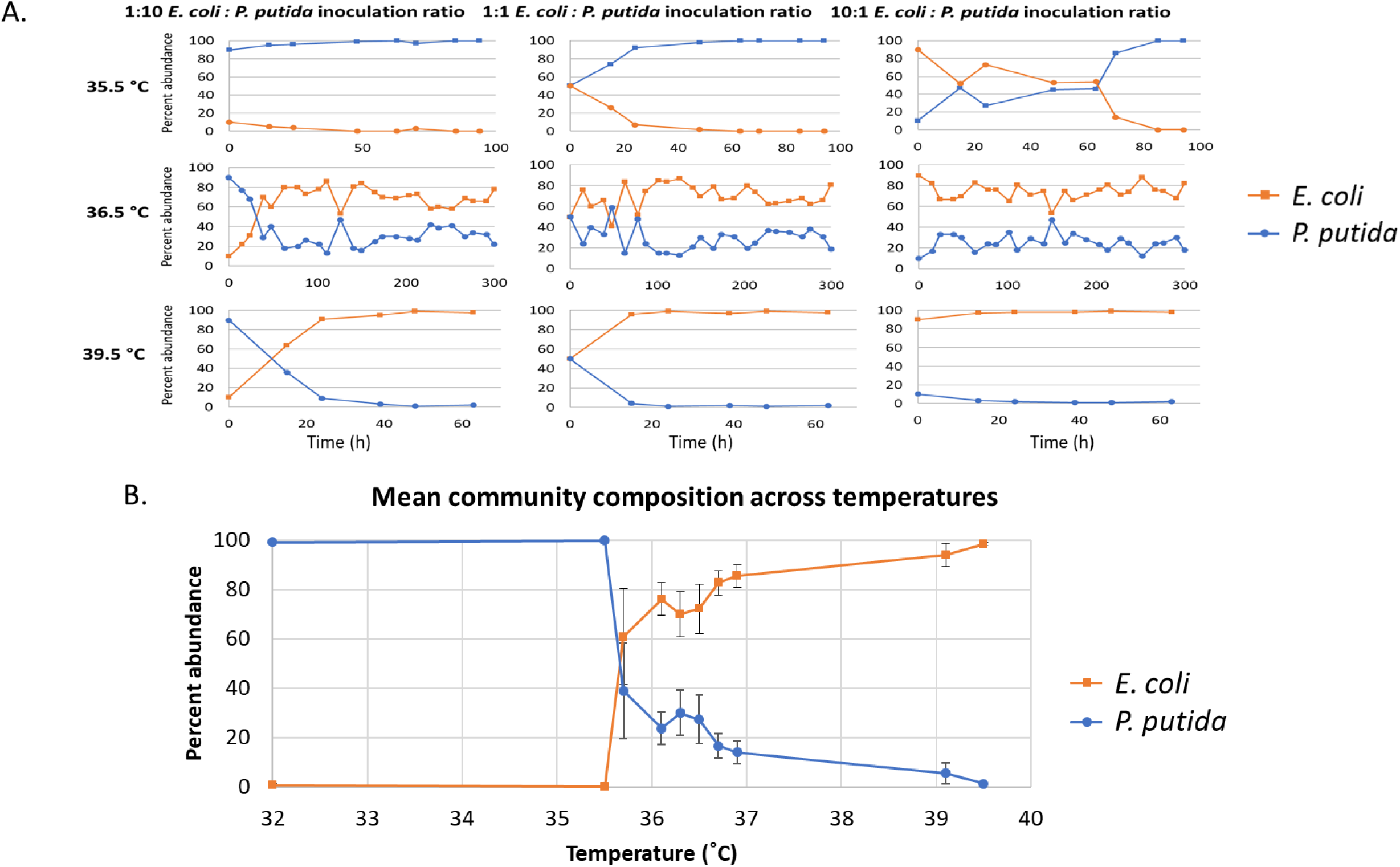
Constant temperature regimes can be defined that result in competitive exclusion of either species as well as coexistence. **A**) Representative flow cytometry data illustrating that when monoculture populations of *E. coli-YFP* and *P. putida-mCh* are identified by front scatter/side scatter (grey boxes) >99% of *E. coli-YFP* exhibit fluorescence under our conditions whereas >99% of *P. putida-mCh* do not (blue boxes). **B**) Representative graphs of species relative abundance over time of bi-culture with varying inoculation ratios grown at different constant temperatures. Each condition has three replicates **C**) Mean community compositions at each temperature.

### 3.4 Regulating community composition using a cycling temperature program

We next explored the effect of temperature on community composition using what we refer to as a cycling temperature regime. In this approach, two distinct temperatures are selected so that the higher temperature favors the growth of one species and the lower one the other species. With these two temperatures selected, the bi-culture is incubated in repeated cycles, of which each starts with growth at the higher temperature for a specific time internal and then switches to growth at the lower temperature for another specific time interval. The key parameters we can manipulate are the two time intervals spent at the two temperatures respectively. When they are selected properly, this scheme enables the two species to coexist. Moreover, the composition of the bi-culture can be tuned as desired and it is determined by a combination of the initial composition and the two time intervals. This can be illustrated analytically if we use a simple mathematical model where each species grows exponentially, which is a reasonable assumption when bacterial cells grow at low densities (i.e. far away from the carrying capacities). Specifically, the two species can be stably maintained provided that the two time intervals are chosen such that the following relationship is satisfied (see Materials and Methods for details):

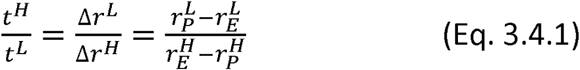

where *t*^*H*^ and *t*^*L*^ are the time intervals the culture spends at the high temperature and low temperature respectively in each cycle. 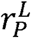 and 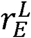 denote the specific grow rates of *P. putida* and *E. coli* respectively at the low temperature. Note that 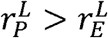 and the difference is defined as 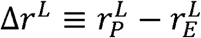. Similarly, 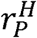 and 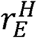 denote the specific grow rates of *P. putida* and *E. coli* respectively at the high temperature. 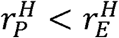 and the difference is defined as 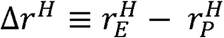. The above equation shows that only the ratio of the two time intervals matters and it is equal to the inverse of the ratio of specific growth rate differences between the two species at the corresponding temperatures. This is a specific quantitative relationship that only holds for the exponential growth model. However, certain qualitative aspects of this relationship are generalizable. In particular, it can be expected that the time interval at which there is a smaller grow rate difference between the two species needs to be longer.

In addition to enabling co-existence, the cycling temperature regime also provides an effective means for tuning the community composition, which is determined by the two time intervals and the initial composition. Taking the simple exponential growth model as an illustrative example again and using the ratio of the two species averaged over a cycle to quantify the composition, we can show that the average ratio over a cycle consisting of *t*^*H*^ at the high temperature and *t*^*L*^ at the low temperature is as follows (see Materials and Methods for details):

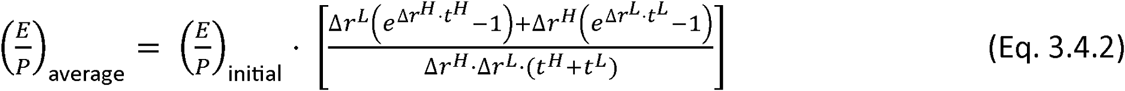

where 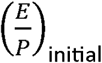 and 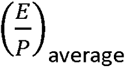 represent the ratio of *E. coli* cells to *P. putida* cells at the beginning and averaged over the cycle respectively. A desired average ratio, therefore, can be achieved when one sets the initial ratio and the two time intervals appropriately. In more general cases where the simple exponential growth model does not apply, the above well-structured analytical relationship may not exist. However, there will still be general trends. For instance, the shorter the two time intervals are, the closer the average ratio is to the initial one. Such design principles can be explored systematically through numerical simulation for specific systems.

Guided by the mathematical modeling and analysis described above, we experimentally implemented the cycling temperature regime on the *E. coli* and *P. putida* bi-culture model system. We were able to demonstrate that the bi-culture could be maintained over an extensive period of time when the parameter settings were chosen appropriately. A representative example is shown in Fig. 4, where 37 °C was chosen as the high temperature favoring *E. coli* and 31 °C as the low temperature favoring *P. putida*. The bi-culture was started with an inoculation ratio of 1:1 and incubated in a cycling temperature program of 16 hours at 37 °C and 8 hours at 31 °C. It was maintained stably at an average composition of 45% *E. coli* and 55% *P. putida* with a standard deviation of 27% for a total of 4 cycles (96 hours).

**Figure 4.**
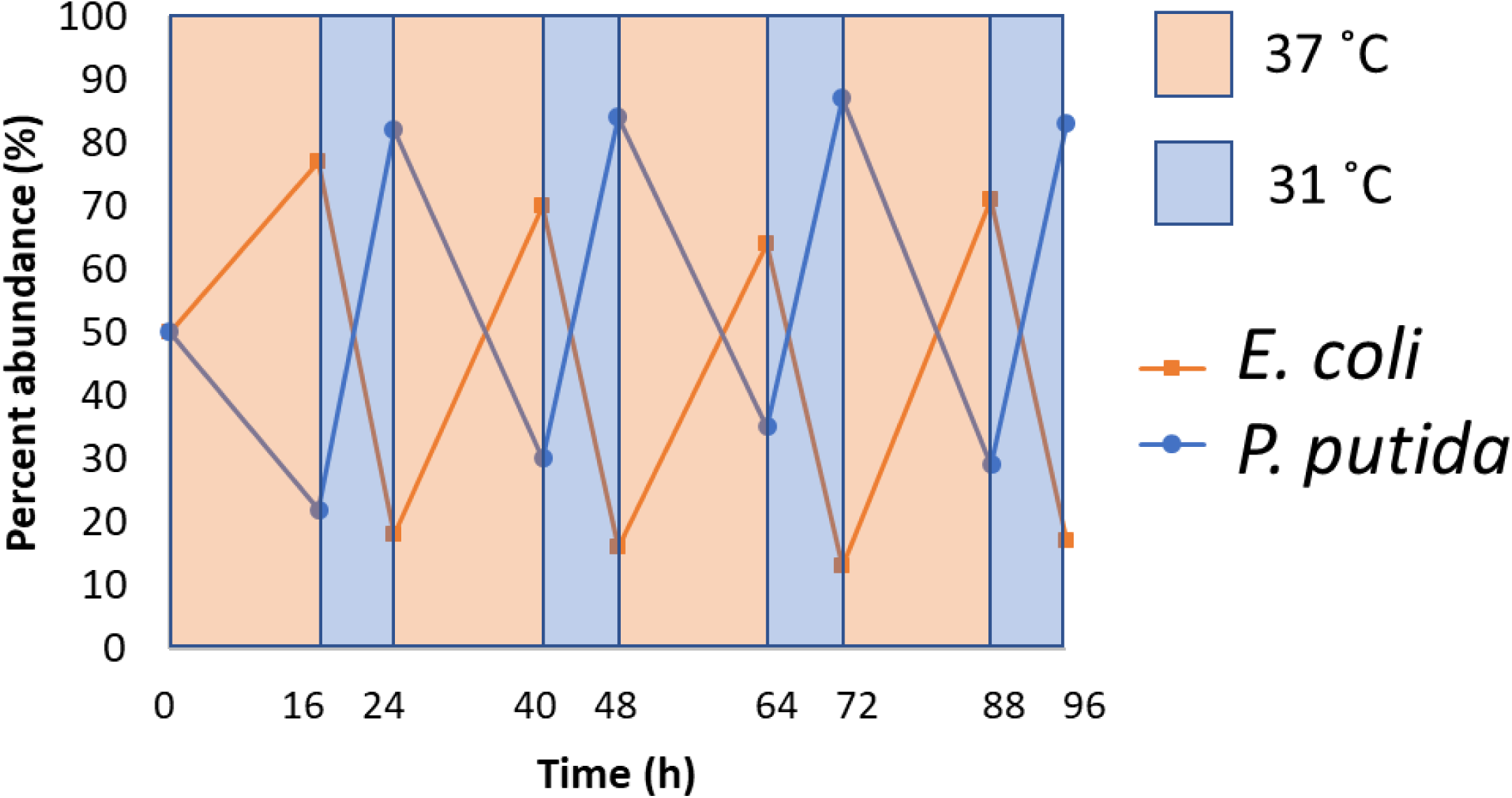
The cycling temperature regime is capable of maintaining coexistence of both species in the bi-culture. With alternating temperatures of which one favors *E. coli* and the other *P. putida*, each species is spared from competitive exclusion.

As illustrated in Fig. 4, the community composition exhibits dynamic oscillations under the cycling temperature regime. The amplitude of the oscillation, i.e. to what extent the composition fluctuates over time, is an important criterion that deserves closer examination. We investigated this first through computational simulation. Here, we assumed no direct interactions between the two species and used two un-coupled logistic growth equations to model the bi-culture incubated in a cycling temperature program and diluted at the end of each time interval (see Materials and Methods for details). Our results showed that the amplitude of the oscillation was dependent on the duration of the time intervals. Longer time intervals result in larger amplitude oscillations in community composition (Fig. 5A) and shorter time intervals lead to more desirable, smaller amplitude oscillations (Fig. 5B). We next validated our model predictions with wet lab experiments. Fig. 6 illustrated the results of an experiment in which we set the temperatures to be 27 °C and 39 °C, which favor *P. putida* and *E. coli* respectively, and incubated the bi-culture with shorter time intervals compared to those in Fig. 4. The bi-culture was maintained at an average composition of 51% *E. coli* and 49% *P. putida* with a standard deviation of 11%, which was substantially smaller than that in Fig. 4, 27%. This approach of employing shorter time intervals can, theoretically, reduce the amplitude of oscillation in community composition to any arbitrarily small value. In practice, however, various constraints would arise. In particular, it takes cells a certain amount of time to physiologically respond to changes in temperature, which sets a lower limit on the time interval parameter in this cycling temperature regime.

**Figure 5.**
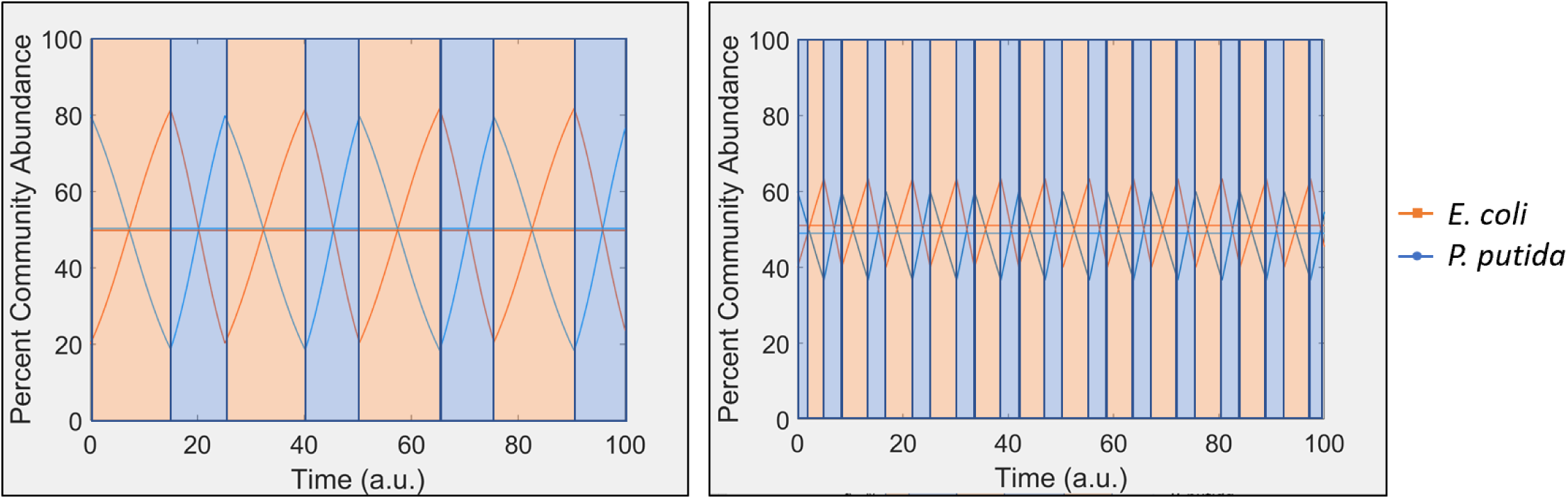
Effect of time intervals on the amplitude of community composition oscillation. Computational simulation shows that when growth related parameters are kept constant, longer time intervals lead to larger amplitude of oscillation (A: *t*^*H*^ = 10.15, *t*^*L*^= 15), whereas shorter time intervals result in more desirable, smaller amplitude of oscillation (B: *t*^*H*^ = 3.4, *t*^*L*^ = 5). In each graph, the horizontal lines represent the average percentages, which are around 50%.

**Figure 6.**
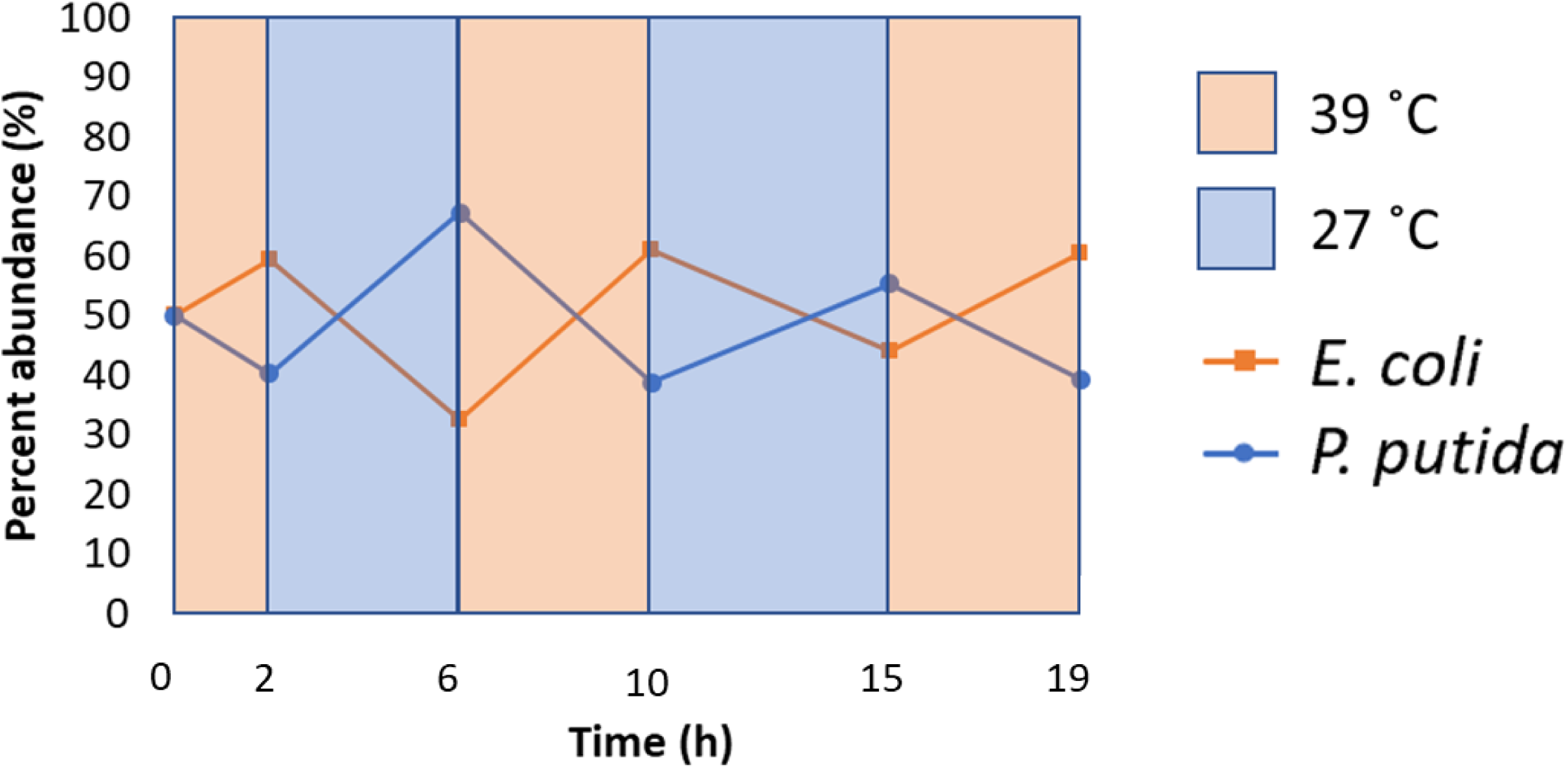
Experimental results demonstrated that shorter time intervals lead to oscillations of smaller amplitudes, in comparison to those in Fig. 4.

## 4. Discussion and Conclusion

Synthetic microbial communities are increasingly being used for a variety of industrial, medical, and environmental applications such as production of commodity chemicals, proteins, and bioremediation (Höffner & Barton, 2014; S. Zhang, Merino, Okamoto, & Gedalanga, 2018). Because microbial community function depends in part on the community composition, it is of inherent interest to synthetic ecology, an emerging area of synthetic biology, that environmental conditions have been demonstrated to affect community composition in an ecological context. While the synthetic biology/ecology literature has yet to treat the subject thoroughly, ecologists have been studying the effects of environmental factors on community composition for decades. It has been proposed and demonstrated that fluctuations in environmental conditions such as nutrient availability and temperature can enable coexistence and influence compositions of various natural communities (Harder & Veldkamp, 1971; Hutchinson, 1961; Jiang & Morin, 2007; Tilman, 1999). Interestingly, in the biotechnology context, the effect of modulating a variety of environmental parameters was explored in the 1970’s and 1980’s (Davison & Stephanopoulos, 1986). These previous studies, however, focused on the chemostat setting, which is not commonly used in biotechnological applications. It is known that an abundance of cellular processes such as nutrient sensing, signal transduction, and macromolecular synthesis are affected by the constantly changing environment within a batch culture (Ziv, Brandt, & Gresham, 2013). Therefore the population dynamics in a batch culture are expected to differ significantly from those seen in a chemostat culture. Accordingly, here we show that in sequential batch cultures, the environmental condition of temperature can be used to modulate growth rates and rationally regulate community composition in synthetic microbial communities.

We show that constant temperature programs can be used to enforce competitive exclusion or to enable co-existence in a synthetic co-culture. It is worth noting that interestingly, the mean community composition does not respond symmetrically to changes in temperature (Fig. 3B). Specifically, we refer to the dramatic shift in community composition from 35.7 °C to 35.5 °C. Above 35.7 °C, the community composition gradually shifts towards higher abundance of *E. coli* as the temperature increases. In drastic contrast, the community composition changes abruptly from primarily *E. coli* to entirely *P. putida* when the temperature decreases by a mere 0.2°C from 35.7°C to 35.5°C. We do not understand the exact cause of this asymmetric behavior, but speculate that interspecies interactions play a role. It is possible that interactions commonly observed between species in microbial co-cultures such as nutrient competition (Ghoul & Mitri, 2016), secretion of toxins (Ghoul & Mitri, 2016), and interspecies crosstalk between quorum sensing systems (Ghoul & Mitri, 2016) occur in our system and could have effects on community composition. It would be interesting for future work to investigate the underlying mechanisms of this intriguing observation.

Control of community composition of synthetic microbial communities is important for optimizing functionality in a biotechnological context. We demonstrate here that temperature regulation is one modality that can be used to rationally regulate composition of synthetic microbial communities. However, a variety of other modalities such as pH, salt concentration, etc. can theoretically be implemented similarly, assuming that the basic requirements we propose for temperature are met (namely that one range of conditions favors growth of one species, and a second range of conditions favors growth of the other species). The advantage of temperature over some of these other modalities is that it can be dynamically changed without requiring media replacement, and is readily implementable. Other modalities for regulating synthetic microbial community composition have been explored in the literature, most notably by Spencer Scott et al. (Scott et al., 2017). In this innovative work, the authors designed and implemented a self-limiting synthetic quorum sensing regulated lysis system to prevent competitive exclusion and enable coexistence between otherwise incompatible community members. One advantage of their approach is the potential for scalability in community complexity, as there are a variety of orthogonal, well-characterized quorum sensing systems that could be utilized with different species or groups of species in a community. Whereas, the alternative approach demonstrated in this work has the advantage that because response to temperature is one of the oldest, core physiological responses of microbes, although adaptation to temperature is expected to diminish its effectiveness at some point, the time scales at which temperature regulation is able to enable coexistence and regulate community composition are quite long (up to 400 hours with our experimental system). However, it would not be straightforward to extend the cycling temperature regime to more complex communities; development of significantly more complicated control schemes will be required.

In this work we have explored temperature as a modality for rationally regulating community composition of a synthetic microbial consortium. While we do not expect the empirical values such as the specific temperatures employed to be applicable to different systems, the conceptual framework along with the associated design principles we have established can be applied to a wide variety of synthetic co-cultures. New approaches for regulating synthetic microbial community composition will continue to emerge and we envision that temperature based control schemes will contribute to a powerful toolbox in the future consisting of various well-developed modalities for regulating synthetic and natural microbial communities, from which researchers can choose based on the specific properties and performance requirements of their system.

## Acknowledgments

AGK was supported by the NIH Cellular and Molecular Biology Training Grant T-32-GM007315 and an Integrated Training in Microbial Systems fellowship. The authors would like to thank Scott Scholz for helpful discussions.

