## Supplementary figures and images for "Temperature regulation as a tool to program synthetic microbial community composition"

### Supplemental figure1

Time series data from all constant temperature regime experiments.

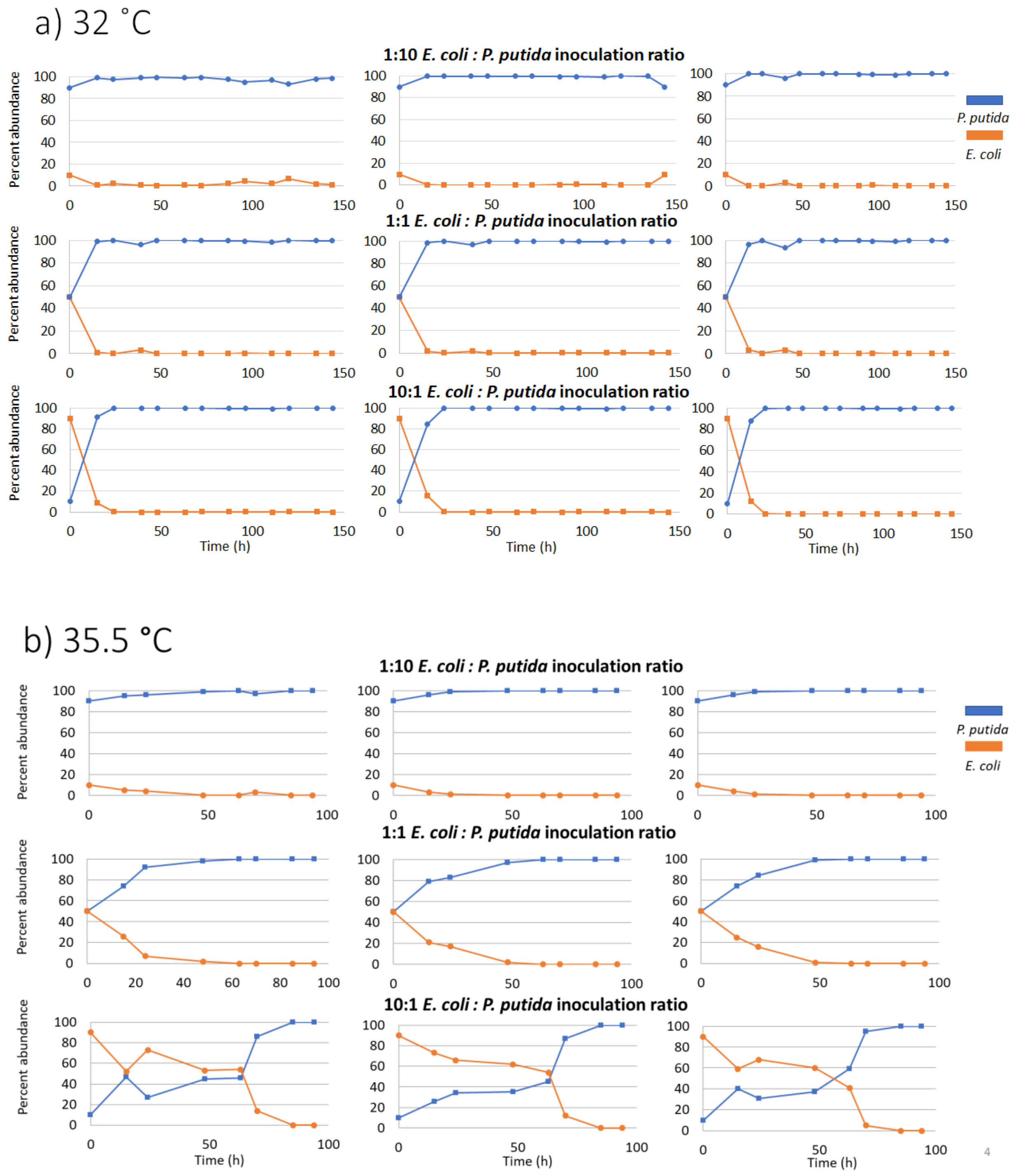

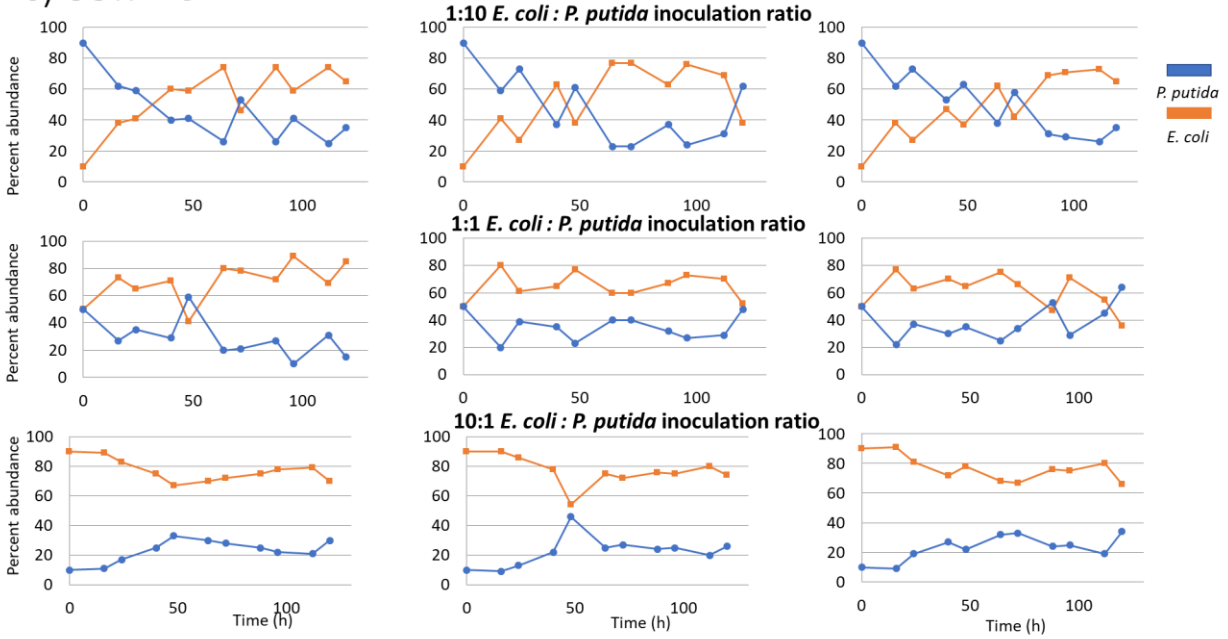

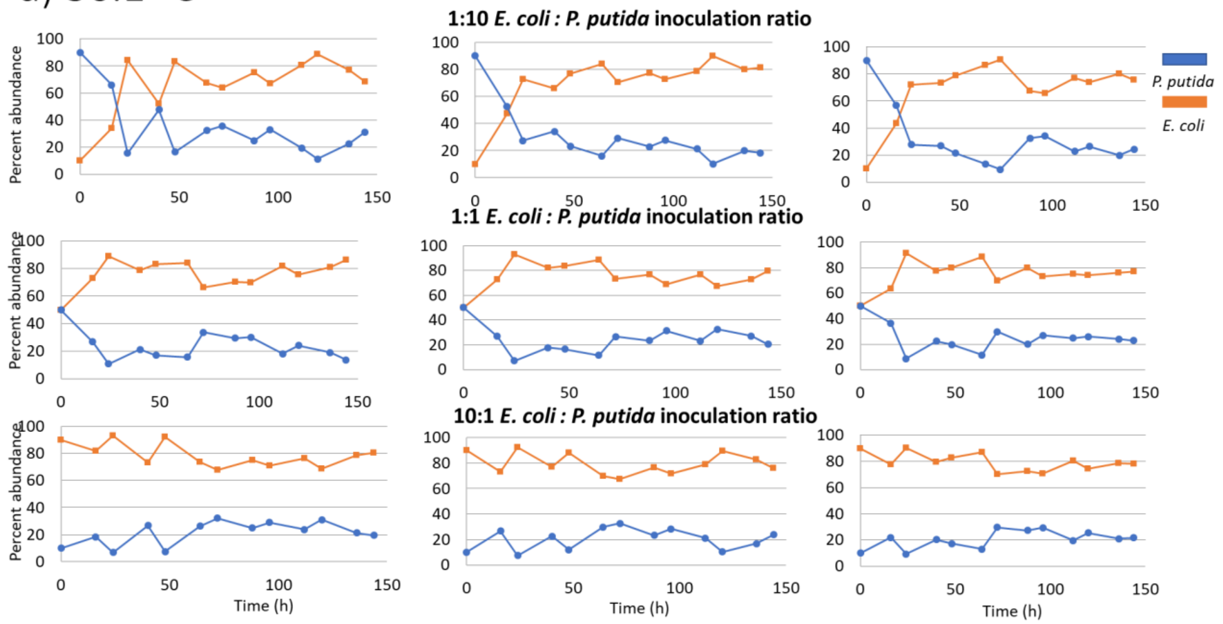

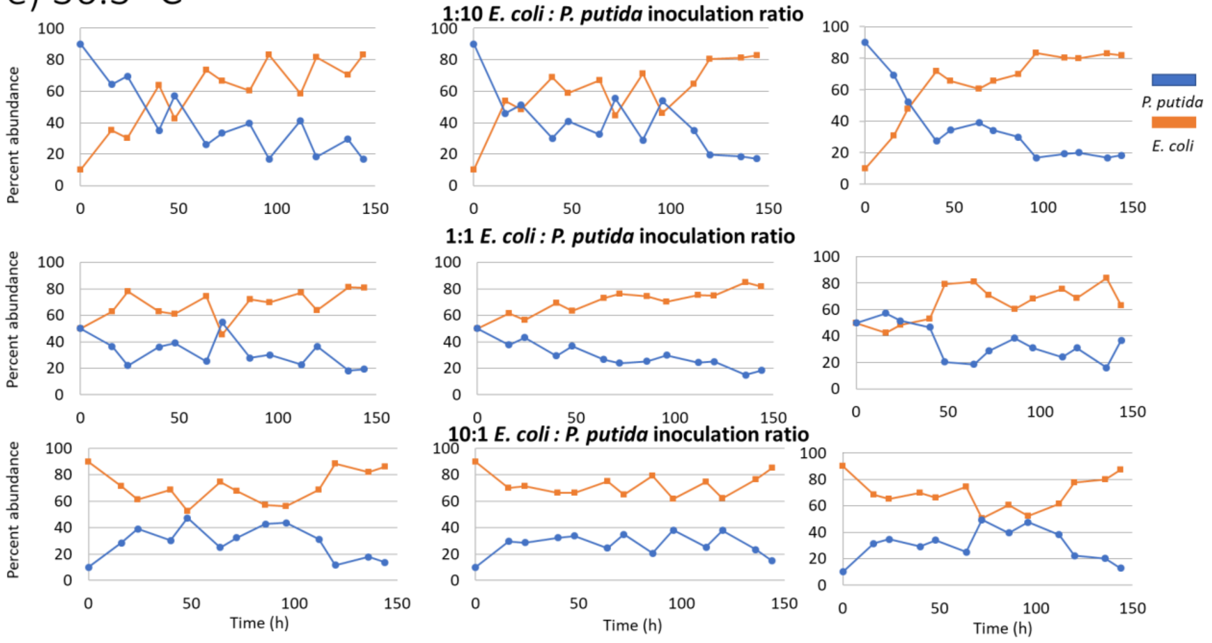

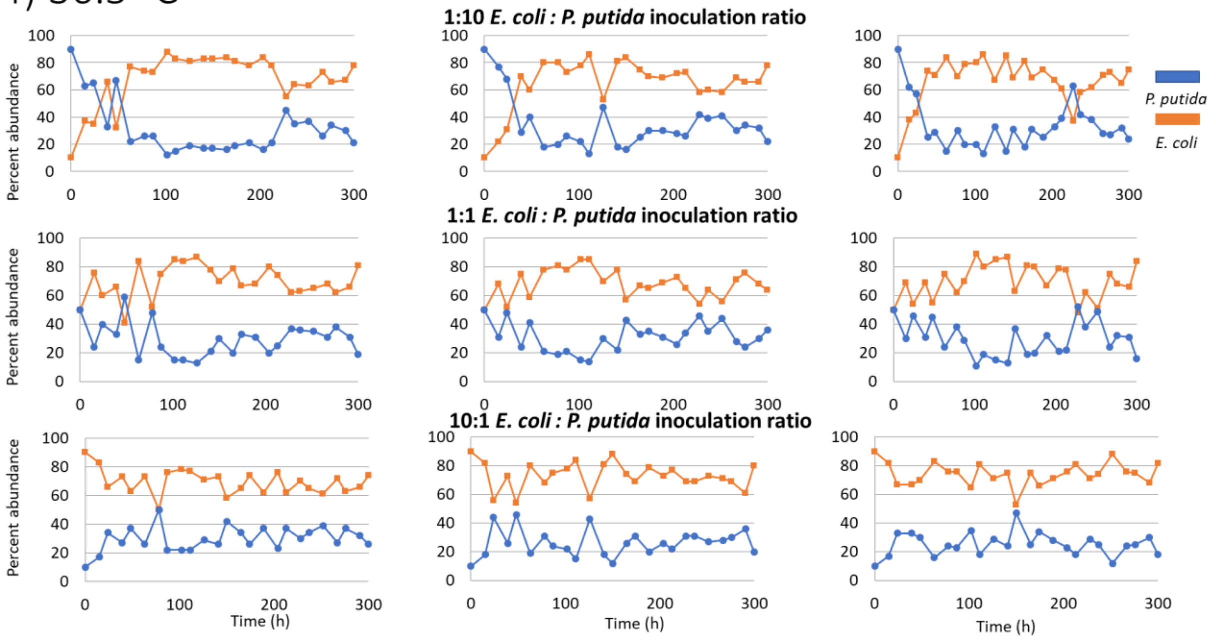

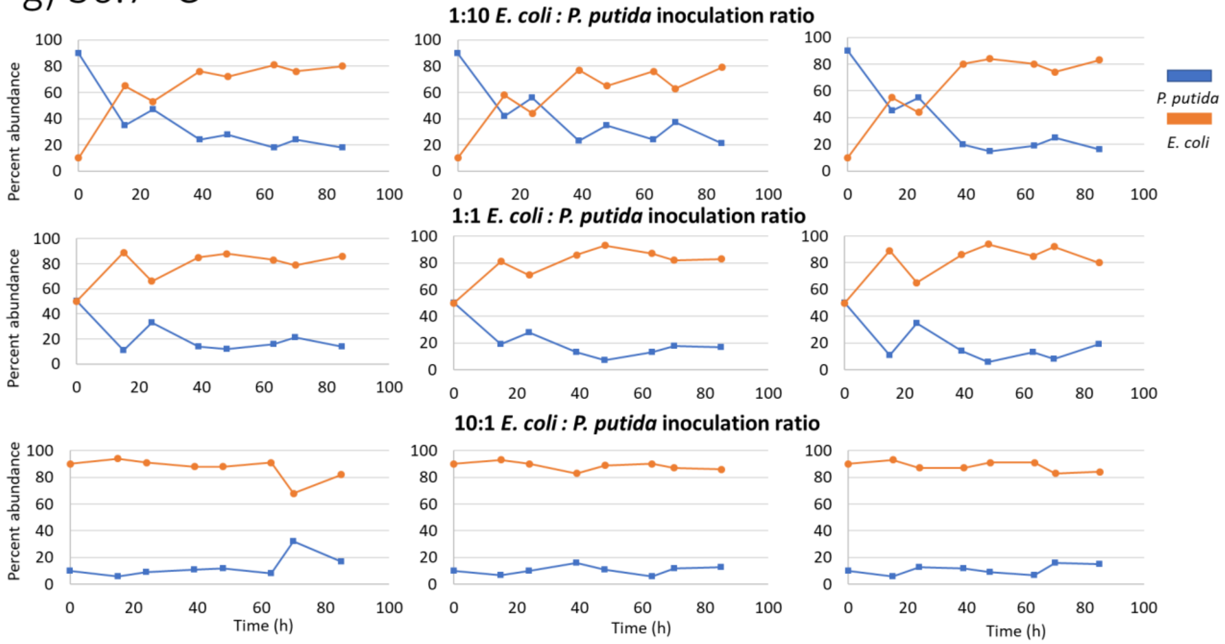

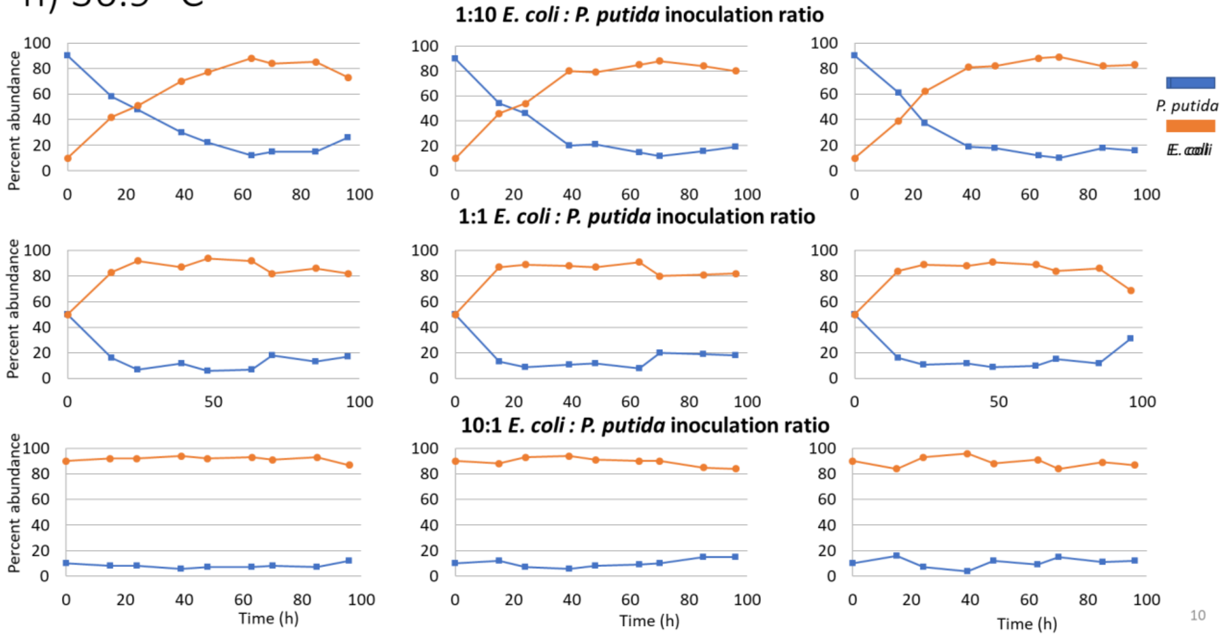

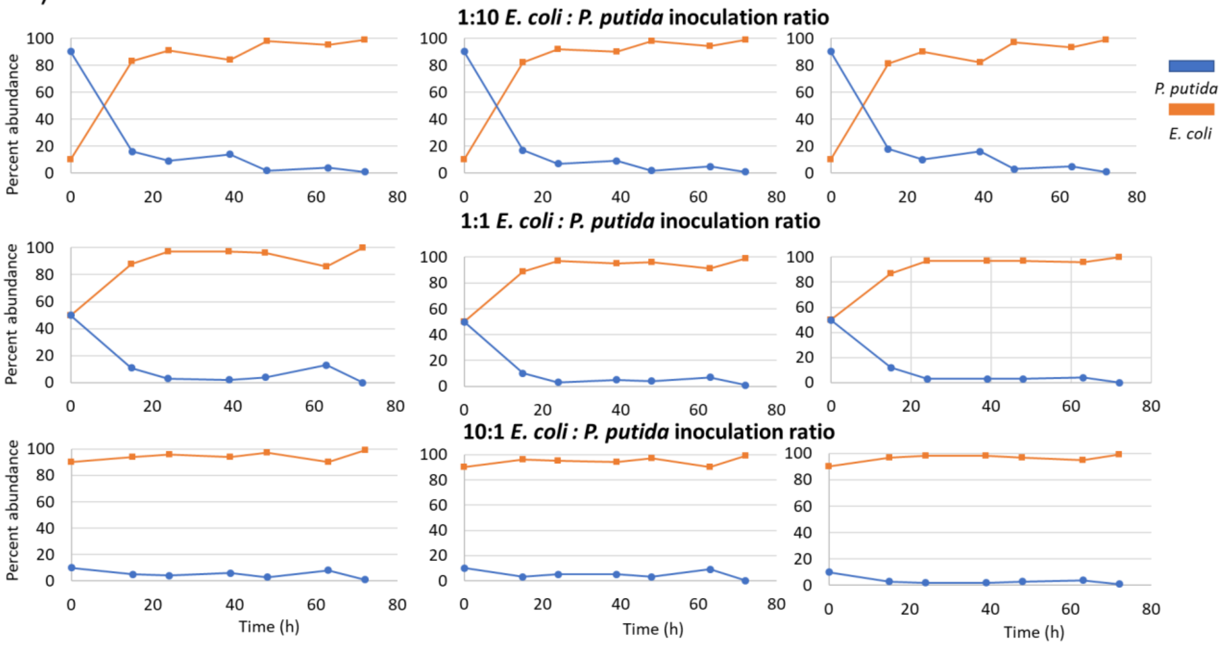

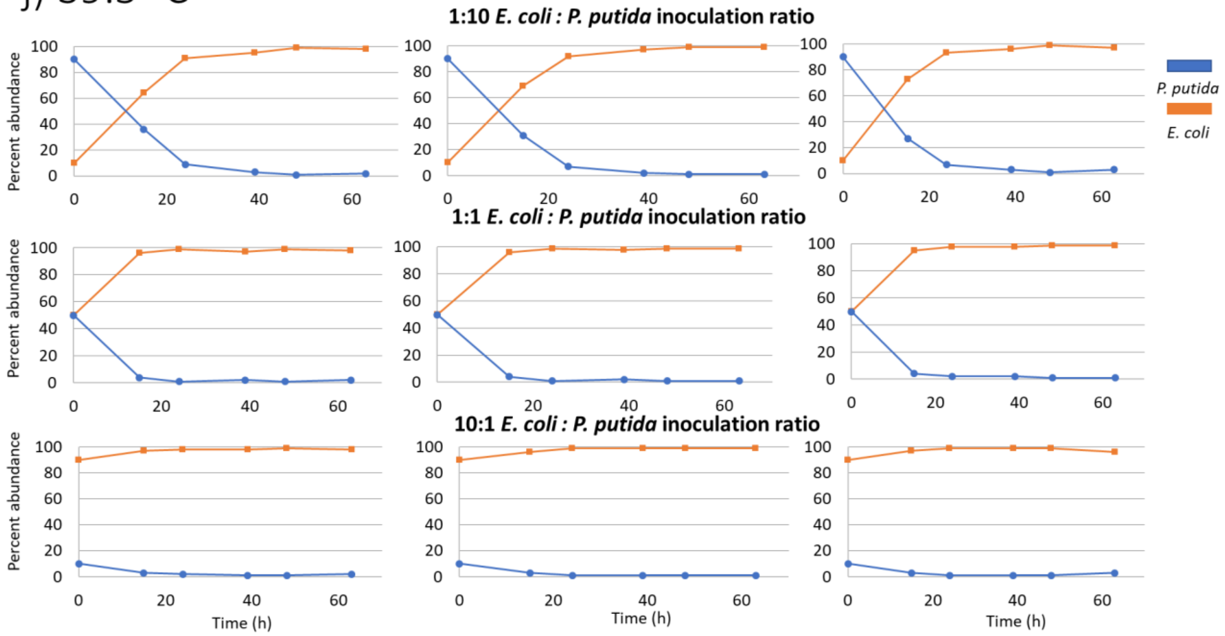
