## Supplemental figure 2 for "Temperature regulation as a tool to program synthetic microbial community composition"

Constitutive expression of a chromosomally integrated cassette for Yellow fluorescent protein (*E. coli*) and mCherry (*P. putida*) were used quantify the community composition of bi-cultures via flow cytometry.

a) b)

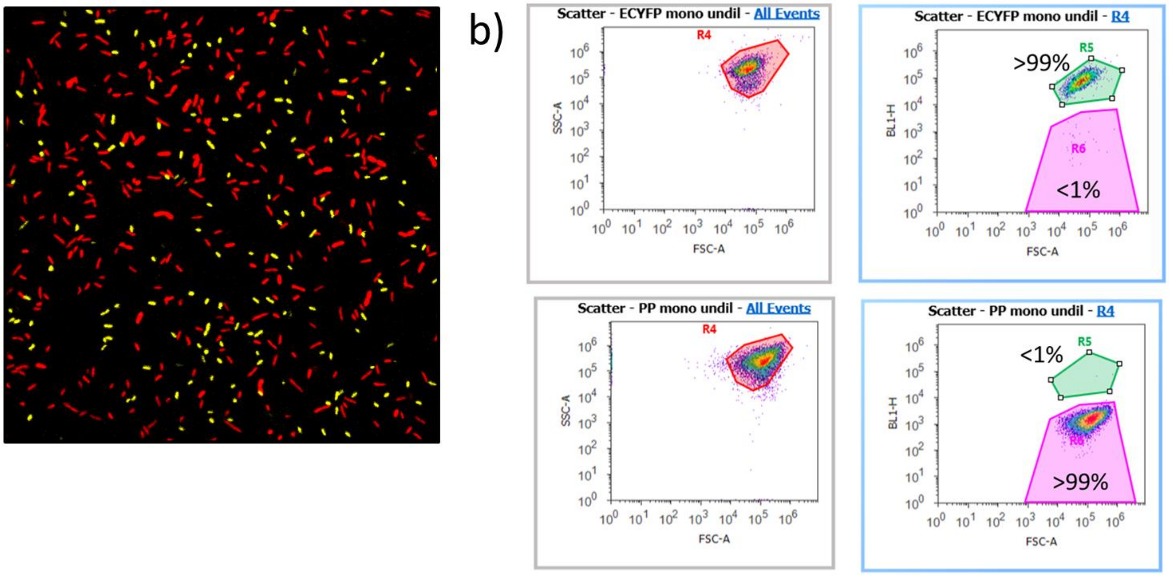
