## Supplemental figure 3 for "Temperature regulation as a tool to program synthetic microbial community composition"

Representative graph of temperatures over time as recorded inside the incubator when the incubator was set to 36.3 ˚C.

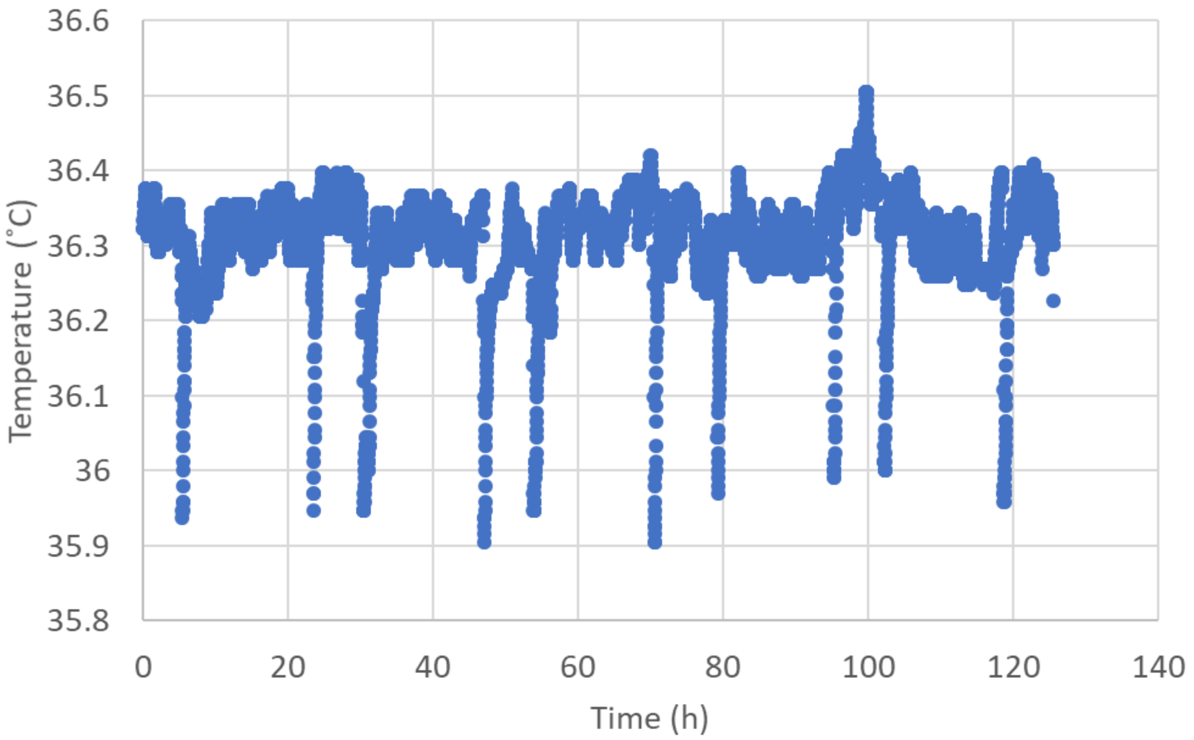
